# EVMP: Enhancing machine learning models for synthetic promoter strength prediction by Extended Vision Mutant Priority framework

**DOI:** 10.1101/2022.10.15.512354

**Authors:** Weiqin Yang, Dexin Li, Ranran Huang

## Abstract

**Motivation:** In metabolic engineering and synthetic biology applications, promoters with appropriate strengths are critical. However, it is time-consuming and laborious to annotate promoter strength by experiments. Nowadays many machine learning (ML) methods are applied to synthetic promoter strength prediction, but existing models are limited by the excessive proximity between synthetic promoters.

**Results:** In order to enhance ML models to better predict the synthetic promoter strength, we propose EVMP(Extended Vision Mutant Priority), a universal framework which utilize mutation information more effectively. In EVMP, synthetic promoters are equivalently transformed into base promoter and corresponding *k*-mer mutations, which are input into BaseEncoder and VarEncoder respectively. In Trc synthetic promoter library, EVMP was applied to multiple ML models and the model effect was enhanced to varying extents, up to 61.30%, while the SOTA(state-of-the-art) record was improved by 15.25%. EVMP also provides optional data augmentation based on multiple base promoters, which further improved the model performance by 17.95% compared with non-EVMP SOTA record. In further study, extended vision is shown to be essential for EVMP. We also found that EVMP can alleviate the over-smoothing phenomenon, which may contributes to its effectiveness. Our work suggests that EVMP can highlight the mutation information of synthetic promoters and significantly improve the prediction accuracy of strength.

**Availability and implementation:** The source code is publicly available on github: https://github.com/Tiny-Snow/EVMP.

**Supplementary information:** Appendix is available at *bioRxiv* online.

## 1. Introduction

Promoters are the fundamental components of transcriptional regulation and have a direct impact on gene expression. In metabolic engineering and synthetic biology applications, promoters with desired strengths are critical (Gao et al., 2021). However, it is challenging for them to meet the requirements of the logical design and optimization of metabolic pathway due to the insufficient number of well-characterized promoters. Therefore, a vast library with hundreds of promoters with better properties must be created and characterized (Tang et al., 2020).

The conventional methods for building promoter libraries rely on functional module combinations or random sequence mutation techniques. Methods based on random sequence mutations, such as error-prone PCR, are regarded as a straightforward and effective mutagenesis technique and have been successfully used to synthesize artificial promoters. Alper et al. (2005), for instance, mutagenized the bacteriophage PL-promoter using error-prone PCR methods. Finally, 22 promoters with an intensity distribution range of 196 times were selected from nearly 200 promoter mutants. Zhao et al. (2021) constructed and characterized a mutant library of Trc promoters (*P*_trc_) using 83 rounds of mutation-construction-screening-characterization engineering cycles, and established a synthetic promoter library that consisted of 3665 different variants, displaying an intensity range of more than two orders of magnitude. Despite the availability of experimental methods, obtaining a small number of useful promoters from a random library typically necessitates time-consuming and laborious screening, and given the enormous number of possible sequence combinations, the effective identification of these useful promoters is largely constrained by a relatively small mutation library. For example, a 20-nucleotide sequence space represents 4^20^ = 1.1 × 10^12^ possibilities, outnumbering even the largest bacterial libraries and most robust high-throughput screens. Therefore, obtaining the desired promoter through mutation, modification, or screening of existing promoters is challenging. In order to effectively guide how to search for new promoters in the huge potential sequence space, new methodologies should be developed to explore the relationship between promoter sequence and function more thoroughly (Cazier and Blazeck, 2021).

Fortunately, prediction of biological problems has been shown to be amenable to machine learning (ML) techniques, as comprehensively reviewed by de Jongh et al. (2020). Several recent studies have applied ML for promoter strength prediction. By varying a 50-nt region in the 5’ UTR of the HIS3 promoter, Cuperus et al. (2017) generated 500000 variants for use as their training dataset for a convolutional neural network (CNN). This allowed them to predictably improve the expression of promoters with both random and native 5’UTRs. They also demonstrated the advantage of using synthetic libraries as they found motifs that enhance transcription that are absent from the yeast genome. Additionally, ML has also enabled the prediction of transcriptional outputs for constitutive, inducible, or synthetic promoters in *Saccharomyces cerevisiae*. For instance, training a CNN with diversified libraries over 10^5^ in size, created by precisely altering the *P*_GPD_ constitutive promoter and a *P*_ZEV_-based inducible system, allowed prediction of promoter strengths with an accuracy of 79% (McIsaac et al., 2014; Kotopka and Smolke, 2020). Similarly, de Boer et al. (2020) created what is perhaps the largest synthetic library ever made for ML promoter engineering by synthesizing over 100 million random 80-bp upstream activating sequences (UASs). This library was used as a training dataset for a TF-motif-based model that was able to correctly predict the expression level of 94% of random promoters. In an elegant combination of hybrid promoter engineering and ML in human cells, Wu et al.(2019) created 129-bp UASs that contained tandem, human TF binding sites that represented over 6000 unique motifs taken from two TF databases, and then used the resulting dataset to train a generalized linear model with elastic net regularization (GLMNET) (Kheradpour and Kellis, 2014; Weirauch et al., 2014; Wu et al., 2019). The model enables us to successfully predict the differential expression of individual promoters across different cell lines, a difficult task for mammalian cell engineering.

Although the progress made in the aforementioned works is exciting, their methods are not without flaws. The key characteristic of the synthetic promoter dataset is that the synthetic promoters are too close to each other while their strengths exhibit a substantial degree of variation (Zhao et al., 2021). This results in the inability to differentiate the strengths of synthetic promoters through sequence homology, and commonly used language models such as LSTM also perform poorly. Therefore, we have considered another simple approach, which is to highlight the mutation information.

In this paper, we propose a novel framework, EVMP (Extended Vision Mutant Priority), to extract important mutation features and enhance ML models. In EVMP, synthetic promoters are equivalently transformed into EVMP format data, including base promoter and *k*-mer mutations (Liu and Wu, 2021), which are input into BaseEncoder and VarEncoder respectively. We evaluated the effectiveness of EVMP on a Trc synthetic promoter library constructed by Zhao et al. (2021) and found that EVMP models exhibited varying degrees of improvement over non-EVMP models. Specifically, LSTM was improved by 61.30%, Transformer by 34.17%, and the other models had effect improvements ranging from 5.15% to 7.27%. Compared to the state-of-the-art (SOTA) non-EVMP method (Zhao et al., 2021), which was included in our experiments, EVMP achieved superior results and improved the SOTA non-EVMP record by 15.25%. EVMP offers optional data augmentation, which involves selecting multiple base promoters and generating multiple copies of data in the EVMP format. Data augmentation further enhanced the effectiveness of EVMP, which was 17.95% higher than that of the SOTA non-EVMP method. We conducted ablation experiments to demonstrate the critical role of the extended vision, or the *k*-mer in mutation representation, in EVMP framework. Additionally, we discussed the effectiveness of EVMP and found that it can alleviate the over-smoothing phenomenon that often occurs in synthetic promoter datasets. Our work suggests that EVMP can significantly improve the performance of ML models in predicting synthetic promoter strength, while its effectiveness is accompanied by reasonable interpretability. To the best of our knowledge, this is the first research to integrate mutation features for predicting promoter strength.

## 2. Methods

### 2.1. EVMP Architecture

As illustrated in Figure 1.A, EVMP framework takes EVMP format data as input and predicts promoter strength using BaseEncoder and VarEncoder as backbone networks. The pipeline for applying the EVMP framework to the synthetic promoter strength prediction task consists of two steps: generating EVMP format data from the synthetic promoter dataset and passing the data through the EVMP framework to obtain prediction results.

**Fig. 1.**
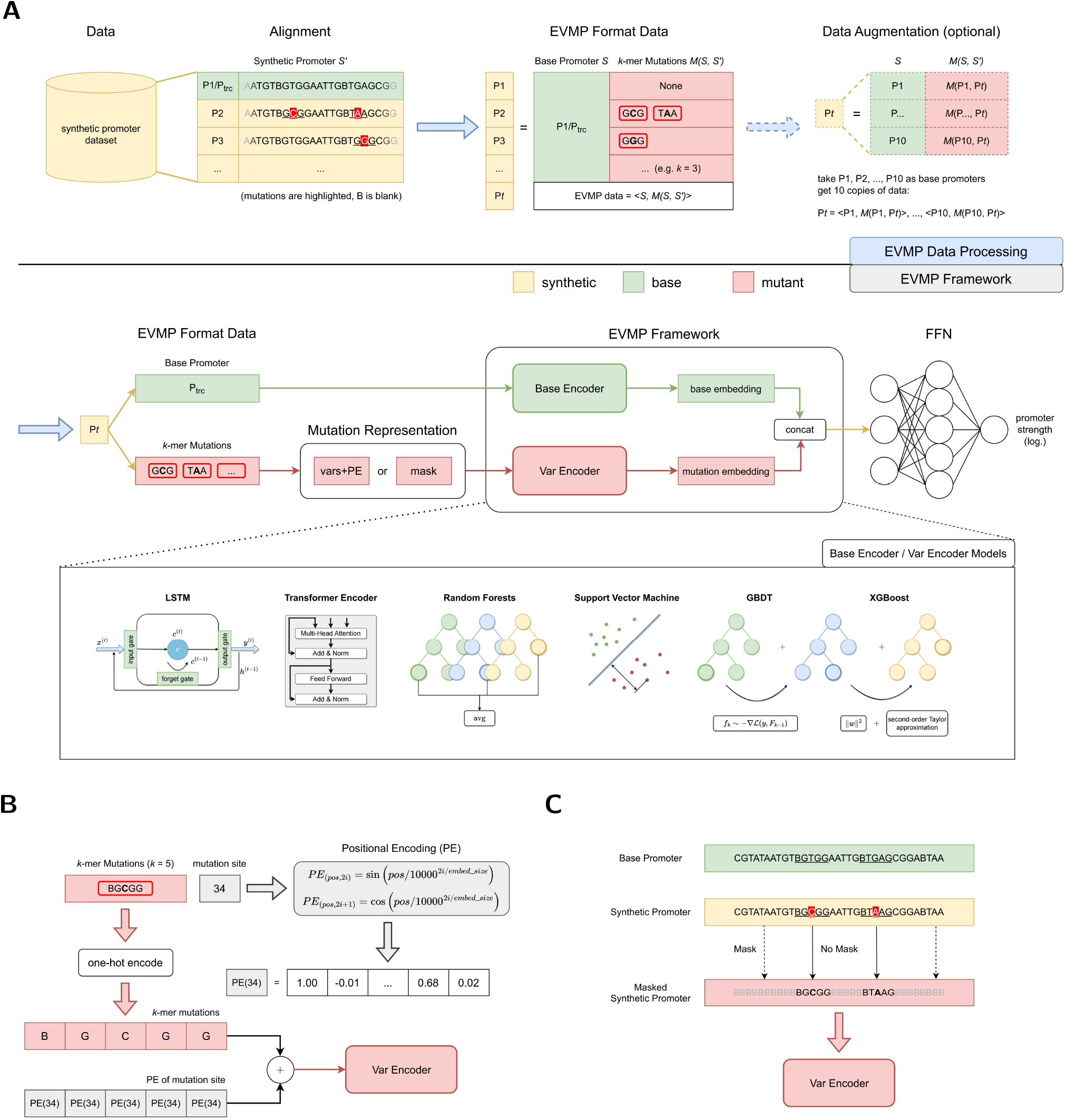
Extended Vision Mutant Priority Framework. **(A)** The overall pipeline of EVMP framework for synthetic promoter strength prediction, which consists of data processing and framework. EVMP data processing is based on alignment, where the original synthetic promoter *S*′ is equivalent to data in EVMP format: the base promoter *S* and the *k*-mer mutation *M*(*S, S*′). *P*_trc_ serves as the default base promoter, while data augmentation based on multiple base promoters is an optional step in data processing. EVMP framework consists of BaseEncoder and VarEncoder, which characterize the base promoter and the *k*-mer mutations after mutation representation, respectively. Base embedding and mutation embedding are then concatenated together and sent to a feed-forward network for strength prediction. **(B)** Vars+PE, one of the mutation representation paradigms. Each base in a *k*-mer mutation is added with the Positional Encoding of the mutation site. **(C)** Mask, one of the mutation representation paradigms. Each base that appears in a particular *k*-mer mutation is retained, and the rest is masked to B (blank).

#### 2.1.1. EVMP Data Processing

In the synthetic promoter dataset, a *base promoter S* = (*x*_1_,…, *x_n_*) is first selected, where each *x* is a base 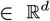 and *d* = 5 (representing one-hot encoding of four bases A, T, C, G, and B for the blank). For any synthetic promoter 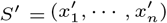, multiple mutation sites are obtained by pairing *S*′ with the base promoter *S*, and the *k*-mer subsequences centered at each mutation site in synthetic promoter *S′* are referred to as *k-mer mutations M*(*S,S′*). Finally, we obtain the EVMP format data consisting of base promoter *S* and *k*-mer mutations *M*(*S, S ′*), which is equivalent to the original synthetic promoter *S*′, denoted as 〈*S,M*(*S, S*′)〉.

#### 2.1.2. Data Augmentation

Data augmentation is an optional step in the EVMP pipeline, which is primarily achieved through selecting multiple base promoters. When data augmentation is not applied, the base promoter is set to *P*_trc_ by default, and each synthetic promoter *S*′ is mapped to a corresponding data point 〈*P*_trc_, *M*(*P*_trc_, *S*′)〉 in the EVMP format. By introducing data augmentation, we can select multiple (e.g., 10 in this experiment) base promoters, denoted as *P*_1_, *P*_2_,…, and generate multiple data points for the same synthetic promoter *S*′. Specifically, we can obtain 〈*P*_1_, *M*(*P*_1_, *S*′)〉, 〈*P*_2_, *M*(*P*_2_, *S*′)〉,…, each corresponding to a different base promoter but the same synthetic promoter. Choosing a suitable base promoter is often challenging. Data augmentation provides the model with more choices and opportunities to select a better base promoter.

#### 2.1.3. EVMP Framework

Similar to common natural language processing (NLP) models, the BaseEncoder in the EVMP pipeline directly processes the base promoter. Specifically, the BaseEncoder takes the base promoter *S* as input and produces the corresponding base embedding *e*_base_ as the output.

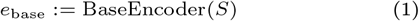

In contrast to typical NLP models that handle the entire synthetic promoter, the VarEncoder in the EVMP pipeline deals with *k*-mer mutations *M*(*S, S*′) between the base promoter *S* and the synthetic promoter *S*′. Specifically, the VarEncoder receives the *k*-mer mutations that are represented by the mutation representation module and produces the corresponding mutation embedding *e*_var_ as the output.

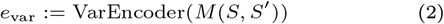

In Section 2.2, we will introduce two different mutation representation paradigms, namely Vars+PE and Mask.

In the EVMP framework, BaseEncoder and VarEncoder are model-agnostic and may not necessarily be distinct. Our study offers several alternatives for BaseEncoder and VarEncoder, including LSTM (Hochreiter and Schmidhuber, 1997; Gers et al., 2000), Transformer (Vaswani et al., 2017), Random Forests(RF) (Breiman, 2001), GBDT (Friedman, 2001), XGBoost (Chen and Guestrin, 2016), and SVM (Cortes and Vapnik, 1995; Boser et al., 1992). For deep learning models, BaseEncoder and VarEncoder can be separate, taking inputs *S* and *M*(*S,S*′), respectively. For conventional machine learning models, BaseEncoder and VarEncoder must be the same model, and the inputs *S* and *M*(*S,S*′) are concatenated. The implementation of the non-EVMP model is the BaseEncoder for each model. In addition, Appendix 1 provides the necessary mathematical foundations for the aforementioned machine learning methods. For more details on model implementations, see Appendix 2.

#### 2.1.4. Promoter Strength Prediction

The base embedding ebase and mutation embedding *e*_var_ produced by the BaseEncoder and VarEncoder, respectively, are concatenated and passed through a feed-forward network (FFN) for predicting promoter strength. The model is trained on a dataset *D* = {(*x_i_, y_i_*)|*i* = 1,…, *N*} using the objective function:

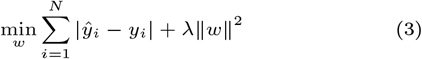

where *ŷ_i_* is the prediction of promoter strength of the EVMP format input *x_i_, w* is the weights of model, and λ is the regularization constant.

### 2.2. Mutation Representation Paradigms

Mutation representation is a critical component of EVMP. Different mutation representation methods may be appropriate for different models. Nevertheless, in general, all mutation representation methods must satisfy two fundamental requirements: (i) the *extended vision* of mutation, and (ii) the preservation of mutation *positional information*. In the scope of our discussion, all mutation representation methods are based on *k*-mer mutations, which aligns with the extended vision principle stated in requirement (i). Therefore, the key to achieving mutation representation lies in preserving positional information. In the following, we present two different paradigms for mutation representation, including Vars+PE (used by LSTM and Transformer) and Mask (used by conventional ML models).

#### 2.2.1. Vars+PE

In Vars+PE, VarEncoder takes the sequence of *k*-mer mutations as input. However, using this form destroys the positional information of individual mutation sites. To incorporate positional information for mutations, one approach is to use Positional Encoding (PE, Vaswani et al. (2017)), which is a function 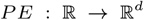. Generally, the Positional Encoding is directly added to the sequence, and therefore, we refer to this technique as *vars+PE*. In this study, we utilize the sin-cos Positional Encoding defined in Definition 1.4 in Appendix 1.4 and Figure 1.B.

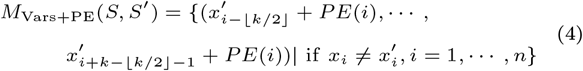

It should be noted that BaseEncoder and VarEncoder add PE in different ways. Specifically, BaseEncoder adds *PE*(*p*) to the base at position *p*, bases at different positions are added different PE, while VarEncoder adds the same *PE*(*p*′) to *k* bases in a *k*-mer mutation at mutated position *p*′, as shown in Equation (4).

#### 2.2.2. Mask

In Section 2.2.1, we utilized the PE technique since the selection of *k*-mer mutations resulted in the disruption of the initial base order. However, if the promoter sequence is maintained in its original form, the provision of additional positional information is unnecessary. The *Mask* technique fits this need, which assigns bases to B(Blank) if they never appear in any of the *k*-mer mutations, so as to disguise the less significant bases.

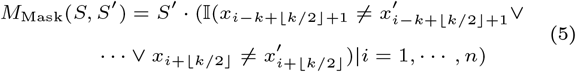

As shown in Figure 1.C, bases that appear in *k*-mer mutations are kept in their original positions, while the remaining bases are masked.

## 3. Experiments

### 3.1. Datasets and Models

The central experiment of this study is the prediction of synthetic promoter strength. The experiments are primarily based on the Trc synthetic promoter library created by Zhao et al. (2021), which used log Fluorescence/OD600 as a measure of promoter strength. Previous research, such as that conducted by Zhao et al. (2021), has evaluated the performance of various models on this dataset, including LSTM, RF, XGBoost and GBDT. Our non-EVMP baselines had included all these competitive models in previous work. In addition, models such as Transformer and SVM were added as options. For EVMP models, BaseEncoder and VarEncoder were the same, and their implementation referred to Section 2.1.3. EVMP is based on alignment, and we used MEGA (Kumar et al., 2008) to perform alignment between the base promoter and synthetic promoter.

Our main experiment used a 9:1 split of the dataset as training data and test data, and the training data was split into a 9:1 training set and validation set in each cross-validation. All experiments used 5-fold cross-validation. It should be noted that our results could not be directly compared to those in Zhao et al. (2021) for fairness since the proportion of the training set was reduced and different independent test sets were adopted. For a fair comparison, the competitive models in Zhao et al. (2021) were all retrained under the same conditions, and we adopted the source code of the original paper, located at https://github.com/YuDengLAB/Predictive-the-correlation-between-promoter-base-and-intensity-through-models-comparing. The results of both EVMP and non-EVMP models for the synthetic promoter strength prediction task were presented in Table 1, and the evaluation metric used is MAE (mean absolute error).

**Table 1.**
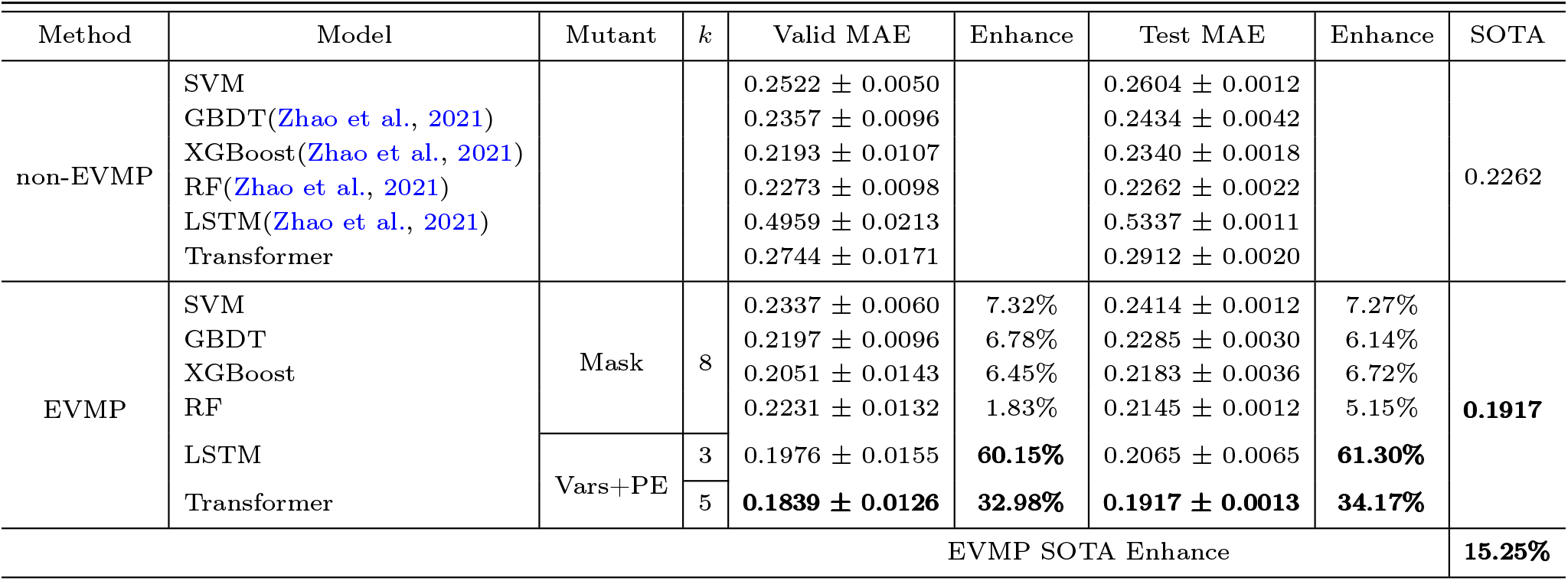
Results of EVMP and non-EVMP models on synthetic promoter strength prediction task. The entry *Mutant* is mutation representation method, *k* is the length of *k*-mer mutations, MAE is mean absolute error, *Enhance* reflects the effect improvement of EVMP models compared to non-EVMP models, measured as the reduction in MAE. All MAEs are the results of 5-fold cross validation and are expressed as mean ± standard deviation.

### 3.2. EVMP Discards Redundant Bases

In the synthetic promoter library, most of the bases between different promoters are identical, which limits the models’ ability to gain informative features from these shared bases. As shown in Figure 2.A, in comparison to *P*_trc_, synthetic promoters with 2, 3 and 4 mutations account for the main part of the dataset, while the vast majority of synthetic promoters have no more than 8 mutations, which indicates that highlighting mutations is necessary.

**Fig. 2.**
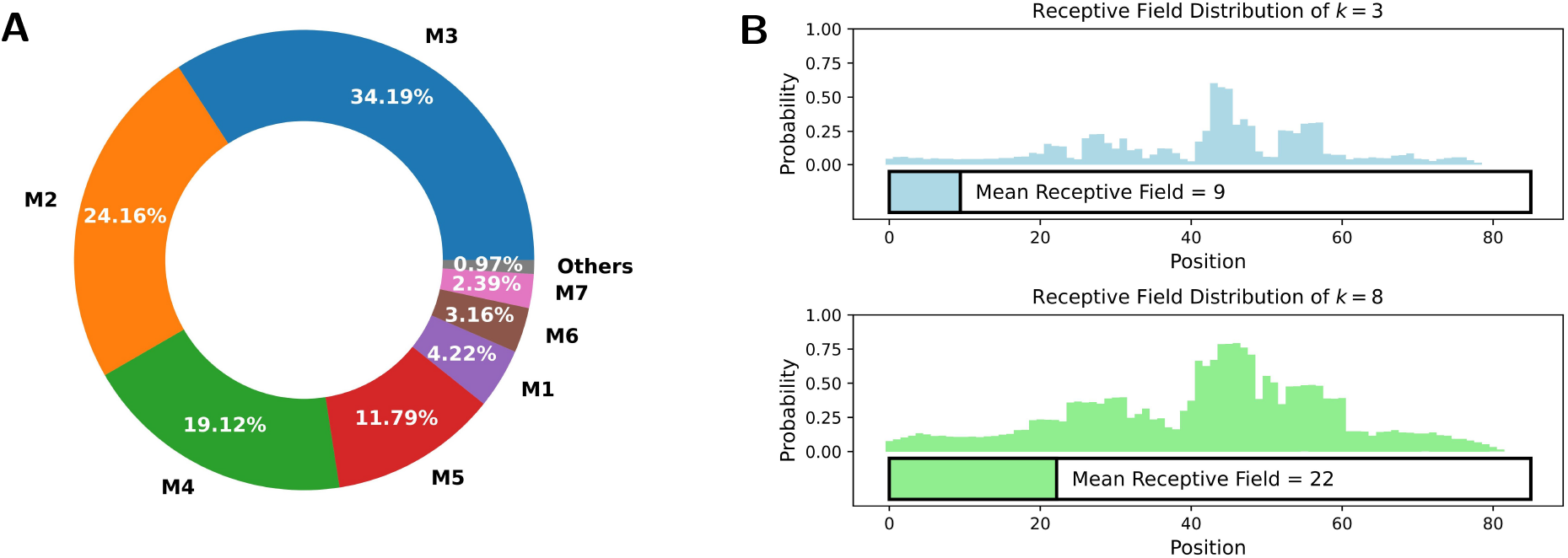
**(A)** The proportion of different mutation times in Trc synthetic promoter library. M*t* (*t* = 1, 2,…) stands for synthetic promoters with *t* different mutation sites. The vast majority (> 99%) of synthetic promoters have no more than 8 mutation sites. **(B)** The probability that each position is included in the receptive field of VarEncoder, each promoter is filled B to 85 length. For *k* = 3 and 8, the mean receptive field of VarEncoder is 9 and 22, respectively, which is much less than the total length of the promoter.

In EVMP, most of the same bases are useless and should be discarded. We define the *receptive field* of VarEncoder as the proportion of the number of bases appearing in a *k*-mer mutation to the length of promoter sequence. As depicted in Figure 2.B, when *k* = 3 and 8, the mean receptive fields of VarEncoder turn out to be 9 and 22 respectively, corresponding to 10.6% and 25.9% of the entire promoter length. The VarEncoder receives inputs that discard redundant bases while retaining necessary mutation information, making the EVMP approach distinct from traditional NLP models.

### 3.3. EVMP Enhances the Effect of ML Models

The results of the comparison between the effects of EVMP and non-EVMP models are presented in Table 1. Among the conventional machine learning models, the state-of-the-art (SOTA) non-EVMP model was RF (0.2262), while the MAE of the other non-EVMP models ranged from 0.23 to 0.26. However, the performance of non-EVMP LSTM (0.5337) and Transformer (0.2912) models was inferior to that of the other ML models.

After applying the EVMP method, all models demonstrated varying degrees of improvement. The LSTM (0.5337 →0.2065) and Transformer (0.2912 → 0.1917) showed a 61.30% and 34.17% improvement, respectively, surpassing the performance of the SOTA non-EVMP model (0.2262) and setting a new SOTA record. Other models also showed improvement ranging from 5.15% to 7.27%. The application of EVMP enhanced the performance of each model in predicting synthetic promoter strength, with the SOTA EVMP model (EVMP-Transformer, 0.1917) outperforming the SOTA non-EVMP model (RF, 0.2262) by 15.25%.

### 3.4. Data Augmentation Further Enhances EVMP

In our previous discussion, we did not incorporate optional data augmentation in the EVMP process. As described in Section 2.1.2, data augmentation involves selecting multiple base promoters (*P*_1_, *P*_2_,…, *P*_10_) and generating multiple copies of data 〈*P*_1_, *M* (*P*_1_, *S*′)〉, 〈*P*_2_, *M*(*P*_2_, *S*′)〉,…, 〈*P*_10_, *M*(*P*_10_, *S*′)〉 for each synthetic promoter *S*′. It is worth noting that the *k*-mer mutations *M* (*P*_1_, *S*′) differ based on the selection of the base promoter *P_i_*, *i* = 1,…, 10. The ten selected base promoters, which differ from *P*_trc_ by 2 to 7 different sites, are listed in Table A4 in Appendix 2.

Table 2 presents the relevant comparative experiments of data augmentation. In these experiments, we chose EVMP-Transformer (*k* = 5) model for analysis. The first two Fixed-Fixed experiments were the same as in Table 1. The third Rand-Rand experiment randomly selected one of the 10 base promoters *P*_1_,…, *P*_10_ as the base promoter, and each synthetic promoter may had a different base promoter. The fourth Augmented-Augmented experiment used all 10 base promoters and expanded the dataset by a factor of 10 by applying data augmentation.

**Table 2.** Results of EVMP-Transformer (*k* = 5) in synthetic promoter strength prediction task with data augmentation. The entries *EVMP* and *DA* indicate whether to use EVMP and whether to use data augmentation, respectively. The entries *Train/Valid* and *Test* refer to the dataset processing method, where *Fixed* refers to using Ptrc as base promoter, *Rand* refers to randomly selecting one of the 10 base promoters *P*_1_,…, *P*_10_ as base promoter, and *Augmented* refers to generating 10 copies of data by using all 10 base promoters. All MAEs are the results of cross validation.

| EVMP | DA | DA Method |  | Valid MAE | Test MAE |
| --- | --- | --- | --- | --- | --- |
|  |  | Train/Valid | Test |  |  |
| × | × | Fixed | Fixed | $0.2744 \pm 0.0171$ | $0.2912 \pm 0.0020$ |
| ✓ | × | Fixed | Fixed | $0.1839 \pm 0.0126$ | $0.1917 \pm 0.0013$ |
| ✓ | × | Rand | Rand | $0.2423 \pm 0.0076$ | $0.2396 \pm 0.0142$ |
| ✓ | ✓ | Augmented | Augmented | **$0.1824 \pm 0.0176$** | **$0.1856 \pm 0.0008$** |
| EVMP+DA SOTA Enhance |  |  |  |  | **17.95%** |

Compared with the Fixed-Fixed method, the EVMP Rand-Rand method (0.2396) exhibited weaker performance than the EVMP Fixed-Fixed method (0.1917), but still outperformed the non-EVMP Fixed-Fixed method (0.2912). It is important to note that the Rand-Rand method represented an extreme scenario, where only approximately 1/10 of the original dataset was utilized by directly using *P*_trc_ as the base promoter. Nevertheless, the Rand-Rand method still achieved better performance than the non-EVMP method, which highlights the robustness of EVMP.

The Augmented-Augmented method with data augmentation (EVMP+DA) achieved the best performance (0.1856) among all experiments, which was 17.95% better than the non-EVMP SOTA. After data augmentation, multiple copies of data in the EVMP format were generated for each synthetic promoter from different perspectives (base promoters). In addition, the Augmented-Fixed method (not listed in Table 2), achieved a MAE of 0.1876 on the same Fixed (*P*_trc_-based) test set in the main experiment, which was better than the EVMP method (0.1917) and lower than the average effect of all base promoters (0.1856). These results suggest that data augmentation indeed increased the diversity of the data to improve the model’s performance, and was able to identify a better base promoter than *P*_trc_. Therefore, data augmentation is a simple and effective alternative to rationally selecting base promoters.

### 3.5. Extended Vision is Necessary for EVMP

Section 2.2 emphasized the crucial importance of extended vision for the effectiveness of EVMP. The inclusion of *k*-mer mutations with *k* > 1 achieves this extended vision. Although it may seem that only including single base mutations (*k* = 1) in the mutation input is the most straightforward approach, a single base may not provide sufficient information for the promoter. As a result, many biological software programs, such as Allesøe et al. (2021) and Nurk et al. (2017), use *k*-mers. Experimental results in Figures 3.A and 3.B, which show the Test MAE of EVMP-Transformer and EVMP-RF for different values of *k* = 1, 3, 5, 8, support this claim.

**Fig. 3.**
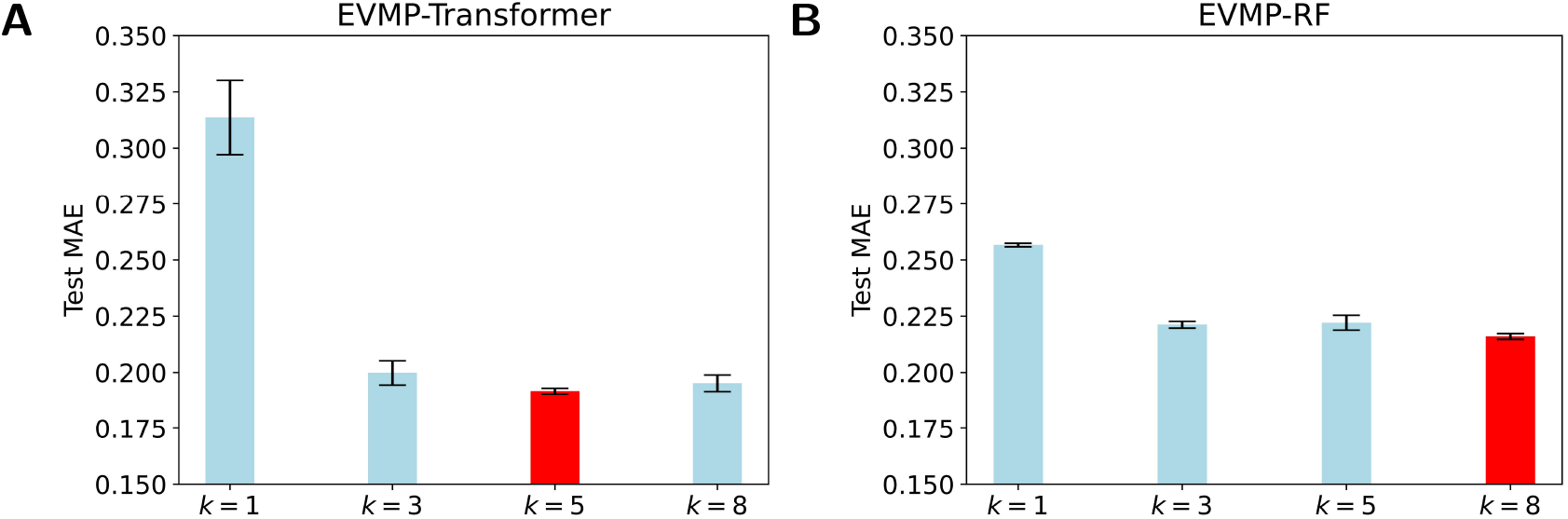
Test MAE of EVMP models with different *k* = 1, 3, 5, 8 (length of *k*-mer Mutations). As *k* increased, the effect of models first improved and then tended to be the same. All MAEs are the results of cross validation. **(A)** Test MAE diagrams of EVMP-Transformer with *k* = 1, 3, 5, 8, 10, *k* = 5 obtained the best result. **(B)** Test MAE diagrams of EVMP-RF, *k* = 8 obtained the best result.

The Test MAE of EVMP-Transformer with *k* = 5 was the best with a value of 0.1917, compared to 0.3136 for *k* = 1, 0.1998 for *k* = 3, 0.1952 for *k* = 8. The Test MAE of EVMP-RF with *k* = 8 was the best with a value of 0.2160, compared to 0.2566 for *k* = 1, 0.2212 for *k* = 3, 0.2221 for *k* = 5. These results indicate that *k* > 1 is much better than *k* = 1 and that extended vision is an indispensable key to the success of EVMP. Furthermore, the effect of the models did not change significantly as *k* increased. Therefore, choosing a suitable *k* is not difficult, and the key is to ensure that *k* > 1. To avoid excessive overlap between *k*-mer mutations, we suggest using the formula *k* = *l*/*α*, where *l* is the length of the promoter and *α* is the average number of mutation sites.

## 4. Discussion

### 4.1. Why EVMP: Experimental Perspective

Experimental results have shown that EVMP can significantly enhance the effect of ML models in predicting the strength of synthetic promoters. However, why EVMP is effective remains to be further explored. We found that EVMP reduces the *over-smoothing* phenomenon, which may contributes to its effectiveness. Over-smoothing phenomenon occurs when the outputs of different inputs appear to be almost the same. Appropriate smoothing is regarded as a symbol of robustness in other tasks like image recognition (Ruderman et al., 2018). However, considering the fact that the differences between synthetic promoters are much shallower, over-smoothing phenomenon is likely to lead to a poor performance.

We define the *average embedding distance* to study oversmoothing phenomenon. Assume that synthetic promoters are *P*_1_,…, *P_N_*, and *P*_0_ is the base promoter (*P*_trc_). The output embedding vector 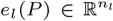 are calculated for each layer *l* and each input *P*. Denote the average embedding distance *d_l_* of each layer as follows:

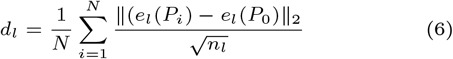

Average embedding distance measures the average distance of each dimension of the embedding at the *l*-th layer, thus it is capable to represent the dispersion of the output of each layer. In particular, a higher average embedding distance indicates a lighter over-smoothing phenomenon.

In Figure 4, the average embedding distance of each layer was calculated for EVMP and non-EVMP implementations of LSTM and Transformer respectively, where LSTM calculated the average embedding distance of each time step. Figure 4.B showed that the average embedding distance of EVMP-Transformer was always much higher than that of non-EVMP Transformer. Figure 4.C showed that the growth rate of the average embedding distance of EVMP-LSTM from 0-th time step was much higher than that of non-EVMP LSTM. The above experimental results demonstrate that EVMP can effectively alleviate the over-smoothing phenomenon.

**Fig. 4.**
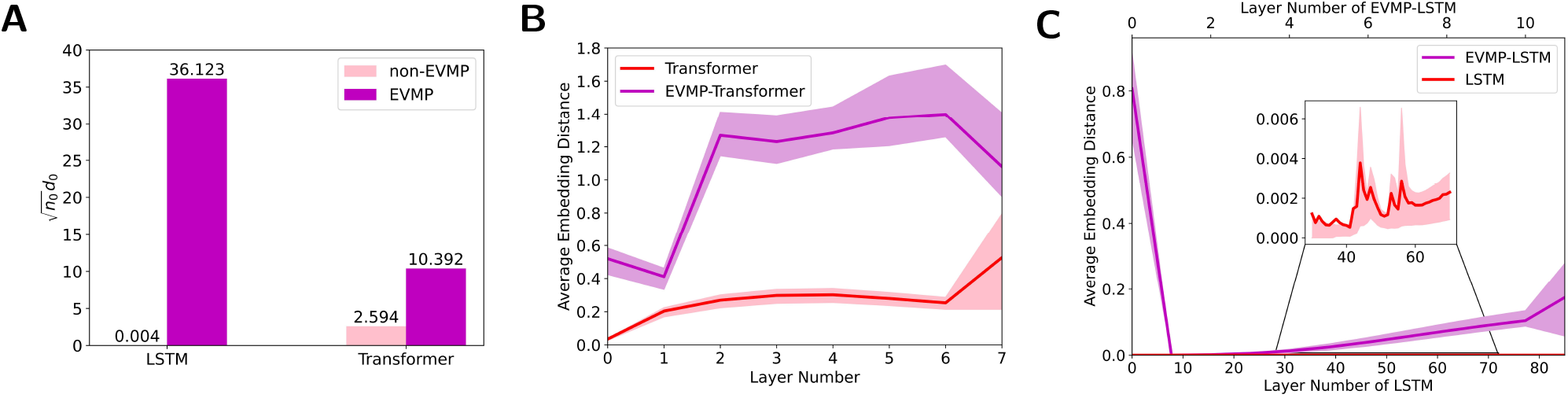
Average embedding distance for each layer of deep learning models, with the 0-th layer serving as the input. Since BaseEncoder receives the same base promoter *P*_trc_, non-EVMP models calculate the average embedding distance of each layer directly, while EVMP models calculate the average embedding distance of VarEncoder. **(A)** non-norm 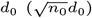 for LSTM and Transformer, which reflects the degree of dispersion of the input data. **(B)** Average embedding distance for each layer of Transformer and EVMP-Transformer. **(C)** Average embedding distance for each layer of LSTM and EVMP-LSTM, where LSTM calculates the average embedding distance for each time step.

### 4.2. Why EVMP: Theoretical Perspective

Section 4.1 offers a comparison of the average embedding distance among layers of neural networks. The purpose is to verify intuitively whether EVMP mitigates the over-smoothing phenomenon and to elucidate the effectiveness of EVMP. In this context, a theoretical illustration is presented to support the validity of this perspective.

Zou et al. (2020) proved that when the width of each layer in deep ReLU networks is at least 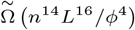, gradient descent can achieve zero training error within 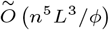 iterations, where *n* is the number of training examples, *L* is the number of hidden layers, and *ϕ* is the maximum lower bound of the *L*_2_-norm between every pair of the training samples.

Here, we regard 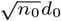 as an approximation of *ϕ*, where *n*_0_ is the dimension of input. As shown in Figure 4.A, EVMP resulted in a noticeable increase in *ϕ* for deep learning models.

Since the size of the training set *n* and the number of model layers *L* are remained unchanged, the overparameterizations of layer width and training iterations are reduced, and the model fitting effect tends to be improved. This statement is also enabled to be applied to RNN and CNN (Zou et al., 2020). The above discussion shows that EVMP can indeed alleviate the over-smoothing phenomenon, which contributes to its effectiveness.

### 4.3. EVMP Reduces the Need for Annotated Data under the Same Prediction Accuracy

Enhancing performance of ML models is just one aspect of EVMP. Another contribution of EVMP is to reduce the requirement for costly strength-annotated synthetic promoters under the original prediction accuracy. Here, given a fixed validation set proportion (1/10 of the original dataset), we randomly selected 1/9, 2/9,…, 9/9 of the remaining 9/10 dataset as training set proportion, respectively. Figure 5 shows MAE and *R^2^* (coefficient of determination) of training and validation sets for each EVMP-Transformer model.

**Fig. 5.**
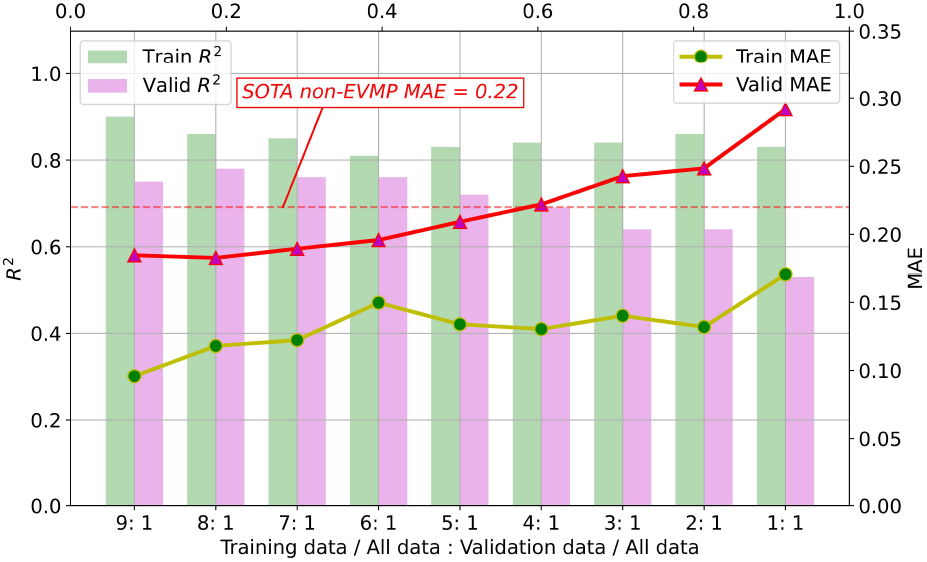
As the proportion of the training set decreased, the MAE of the validation set increased and the *R^2^* decreased. When the proportion of training set to validation set is higher than 5:1, EVMP-Transformer achieved lower MAE than SOTA non-EVMP models.

According to our experimental results in Table 1, the SOTA non-EVMP MAE was approximately 0.22, while Figure 5 shows that EVMP only needed a training set of 5/9 original size to achieve this effect. In other words, EVMP only needed 56% of the original data while maintaining the same prediction accuracy, which greatly reduces the dependence on high-quality labeled data.

## 5. Conclusion

In this work, we proposed EVMP as a universal framework to enhance machine learning models for synthetic promoter strength prediction, which includes two steps: EVMP data processing and EVMP framework. The original synthetic promoter data are transformed into base promoter and *k*-mer mutations, which are processed by BaseEncoder and VarEncoder respectively to highlight mutation information. EVMP enhanced many models and achieved new SOTA records. EVMP also provides optional data augmentation based on multiple base promoters, which can further improve the performance of EVMP. In further study, We experimentally verified that extended vision, or *k*-mer, is critical for the effectiveness of EVMP. In terms of interpretability, EVMP is proved to be able to alleviate the over-smoothing phenomenon and thus improves the effect of models.

EVMP is a highly robust and versatile machine learning enhancement framework with the potential to be extended to various mutation-based synthetic biology component libraries. In future research, other ways of mutation representation and more efficient information transfer between BaseEncoder and VarEncoder can be further explored.

## Supporting information

EVMP_Appendix

## Supplementary Material

Appendix is available at https://www.biorxiv.org.

## Data Availability

The source code is available at https://github.com/Tiny-Snow/EVMP, including all non-EVMP and EVMP models implemented in the paper.

