## Supplementary material for "EVMP: Enhancing machine learning models for synthetic promoter strength prediction by Extended Vision Mutant Priority framework": EVMP_Appendix

### 1. Supplement on Machine Learning

#### 1.1. Decision Tree Models

**Definition 1.1** *A CART (classification and regression tree) is*

$$f(x) = \sum_{m=1}^{M} c_m \mathbb{I}_{R_m}(x) \tag{A1}$$

*where $R_m(m = 1, \cdots, M)$ is a partition of the input space, and $c_m$ is the unique output of all input $x \in R_m$.*

Suppose the training dataset is $D = \{(x_i, y_i)|i = 1, \cdots, N\}$ , we use square error to represent the error of CART to training data. From derivation, it's easy to prove that

**Theorem 1.1** *The optimal value of $c_m$ is*

$$\hat{c}_m = \text{avg}(y_i|x_i \in R_m) \tag{A2}$$

**Definition 1.2** *The least squares regression tree is a CART splitting algorithm that generates CART. For an input space $\mathcal{R}$, a splitting refers to finding a dimension $j$ of input $x$, selecting a splitting point $s$ on the dimension $j$, and dividing $\mathcal{R}$ into two regions*

$$\mathcal{R}_1 = \{x|x^{(j)} \leq s\}, \mathcal{R}_2 = \{x|x^{(j)} > s\} \tag{A3}$$

Theorem 1.1 gives the optimal solution $c_1, c_2$ to the square error $L = \sum_{x_i \in \mathcal{R}_1}(y_i - c_1)^2 + \sum_{x_i \in \mathcal{R}_2}(y_i - c_2)^2$. The global optimal solution is given by traversing $j$ and $s$, and it is easy to prove that $s$ must be equal to one of $x_i^{(j)}(i = 1, \cdots, N)$.

Random Forests (Breiman et al., 1984) are *bagging* tree model based on CART, while GBDT (Gradient Boosting Decision Tree, Friedman (2001)) and XGBoost (eXtreme Gradient Boosting, Chen and Guestrin (2016)) are *boosting* tree model. In the following discussion, we only use the properties of CART.

#### 1.2. Support Vector Machine (SVM)

Support Vector Machine (SVM) is a linear classifier (Cortes and Vapnik, 1995). In the following discussion, we omit the kernel trick (Boser et al., 1992).

In essence, for a dataset $D = \{(x_i, y_i)|i = 1, \cdots, N\}$, the purpose of SVM is to find a hyperplane $w \cdot x + b = 0$, with the functional distance $\hat{y}_i = w \cdot x_i + b$ between any input vector $x$ and the hyperplane as the regression predictive value.

**Definition 1.3** *Linear SVM regressor is a optimization problem as follows*

$$\min_{w,b} \frac{1}{2}\|w\|^2 + C\sum_{i=1}^{N} L_\varepsilon(\hat{y}_i, y_i) \tag{A4}$$

*where $C > 0$ is a penalty parameter, $L_\varepsilon$ is $\varepsilon-$Insensitive error function* (Theodoridis, 2015)

$$L_\varepsilon(\hat{y}_i, y_i) = \begin{cases} 0, & \text{if } |\hat{y}_i - y_i| < \varepsilon \\ |\hat{y}_i - y_i| - \varepsilon, & \text{otherwise} \end{cases} \tag{A5}$$

A few tricks can be used to turn Problem (A4) into a convex optimization problem.

$$L_\varepsilon(\hat{y}_i, y_i) = \xi_i + \xi_i' \tag{A6}$$

where $\xi_i$ and $\xi_i'$ satisfy

$$\begin{aligned} \xi_i \geq 0 \quad & y_i - \hat{y}_i - \xi_i \leq \varepsilon \\ \xi_i' \geq 0 \quad & \hat{y}_i - y_i - \xi_i' \leq \varepsilon \end{aligned} \tag{A7}$$

Obviously, Problem (A4) remains the same after the substitution of Equation (A6) and (A7). For the new convex optimization problem, it is easy to get the optimal solution (A9) by Lagrange multiplier method (A8).

$$\begin{aligned} \max_{\alpha,\alpha',\beta,\beta'} \min_{w,b,\xi,\xi'} L = & \frac{1}{2}\|w\|^2 + C\sum_{i=1}^{N}(\xi_i + \xi_i') \\ & + \sum_{i=1}^{N}\alpha_i(y_i - \hat{y}_i - \xi_i - \varepsilon) + \sum_{i=1}^{N}\alpha_i'(\hat{y}_i - y_i - \xi_i' - \varepsilon) - \sum_{i=1}^{N}\beta_i\xi_i - \sum_{i=1}^{N}\beta_i'\xi_i' \end{aligned} \tag{A8}$$

The optimal solution is (omit other results)

$$w^* = \sum_{i=1}^{N} (\alpha_i - \alpha_i') x_i \tag{A9}$$

Substituting into the optimal solution, we obtain

$$\begin{aligned} & \min_{\alpha,\alpha'} \frac{1}{2} \left( \sum_{i=1}^{N} (\alpha_i - \alpha_i') x_i \right)^2 + \sum_{i=1}^{N} (\alpha_i + \alpha_i') y_i + \varepsilon \sum_{i=1}^{N} (\alpha_i + \alpha_i') \\ & \text{s.t. } 0 \le \alpha_i, \alpha_i' \le C, i = 1, \cdots, N \end{aligned} \tag{A10}$$

Problem (A10) is what we should solve by dataset $D$.

#### 1.3. Mask EVMP in Machine Learning Models

In this appendix, we briefly analyze the properties of mask EVMP ($k = 1$). Although EVMP's effectiveness depends on the specific problem and dataset distribution, we can still find some useful properties of mask in machine learning models, including SVM and CART.

Mask is a technique that, in dataset $D = \{(x_i, y_i) | i = 0, \cdots, N, x_i \in \mathbb{R}^n\}$, we select a vector $x_0$, for the remaining vector $x_i (i = 1, \cdots, N)$, the masked vector is $x_i'$ s.t. $\forall j = 1, \cdots, n$

$$x_i'^{(j)} = \begin{cases} x_i^{(j)}, & \text{if } x_i^{(j)} \neq x_0^{(j)} \\ 0, & \text{otherwise} \end{cases} \tag{A11}$$

If $x$ is a concatenation of multiple one-hot vectors, or more generally, if $x$ is equal to 0 or 1 in all dimensions, we observe the following properties

**Lemma 1.1** *Mask is a technique to remove dimensions, which is reflected in the one masked dimension will all become zero.*

$$\left( \begin{array}{cc|c|ccc} B: & 1 & 0 & 0 & 0 & 0 \\ \hline A: & 0 & 1 & 0 & 0 & 0 \\ \hline G: & 0 & 0 & 1 & 0 & 0 \\ C: & 0 & 0 & 0 & 1 & 0 \\ T: & 0 & 0 & 0 & 0 & 1 \end{array} \right) \Longrightarrow \left( \begin{array}{c|c|ccc} 1 & 0 & & & \\ & 0 & & & \\ & 0 & 1 & & \\ & 0 & & 1 & \\ & 0 & & & 1 \end{array} \right) \tag{A12}$$

This property is easily observed in Equation (A12). The mask technique sets some specified dimensions within the one-hot encoding to zero. Therefore, Lemma 1.1 guarantees that some dimensions are the same in all $x_i$, so we can apply Lemma 1.2.

**Lemma 1.2** *If some dimensions are the same for all $x_i$, then the solutions of SVM and CART remain the same after removing these dimensions.*

*Proof* Given the symmetries between the dimensions, suppose $x_i = [x_i^{(1)}, \cdots, x_i^{(m)}, c^{(1)}, \cdots, c^{(n-m)}]$, where $c$ is the same dimensions of all $x_i$.

First to prove that the solution of SVM remains the same. From Equation (A8) and $\partial L / \partial b = 0$, we obtain

$$\sum_{i=1}^{N} \alpha_i = \sum_{i=1}^{N} \alpha_i' \tag{A13}$$

By Equations (A9) and (A13), we have

$$w_1^* = \left[ \sum_{i=1}^{N} (\alpha_{1i} - \alpha_{1i}') x_i^{(j)} \right]_{j=1,\cdots,m}, \quad w_2^* = \left[ \sum_{i=1}^{N} (\alpha_{1i} - \alpha_{1i}') x_i^{(j)}, \mathbf{0}^{n-m} \right]_{j=1,\cdots,m} \tag{A14}$$

Since optimization problems (A10) on for $w_1^*$ and $w_2^*$ are equivalent, the solutions of SVM are the same.

Second, we show that the results of CART also remain the same. By Definition 1.2, CART keeps the same optimal splitting points in the dimensions $x_i^{(1)}, \cdots, x_i^{(m)}$. In the dimension $c^{(1)}, \cdots, c^{(n-m)}$, according to CART splitting algorithm in Equation (A3), since $\forall x_i$ have the same values in these dimensions, nodes will not be split in these dimensions. Therefore, the CARTs generated by $D = \{(x_i, y_i) | i = 0, \cdots, N\}$ and $D' = \{(x_i \backslash c, y_i) | i = 0, \cdots, N\}$ are the same, and the results are also the same guaranteed by Theorem 1.1. □

From Lemmas 1.1 and 1.2, we know that once the mask technique is used, some dimensions are equivalent to being removed. These dimensions are what are called redundant features. A typical example is the CART generated by five one-hot sequences in Equation (A12). Even if one feature is removed, it can be obtained from other features. Removing these redundant features will make CART not split in unnecessary dimensions and avoid strong over-fitting of CART. On the other hand, because of regular terms and depth constraints in common decision tree models, mask will make CART get better classification results under the same depth constraints and regularity penalties. Therefore, we have the following corollary:

**Corollary 1.1** *Compared with non-mask technique, for single CART with mask technique, the same results can be obtained; for CART models with regular terms and depth constraints, the same or better results can be obtained using the mask technique.*

The effect of mask technique on regular terms is significant. We show that mask technique can reduce the upper bound of regular terms in SVM.

**Lemma 1.3** *The mask technique reduces the upper bound of the regular term* $\frac{1}{2}\|w\|^2$ *in SVM.*

*Proof* By Karush–Kuhn–Tucker optimality conditions (Kuhn and Tucker, 2014), we obtain

$$\alpha_i \alpha_i' = 0, \text{ for } i = 1, \cdots, N \tag{A15}$$

That means at least one of $a_i$ and $a_i'$ is 0. So

$$|\alpha_i - \alpha_i'| \leq |\alpha_i| + |\alpha_i'| \leq C \tag{A16}$$

Therefore, we obtain

$$\frac{1}{2}\|w\|^2 \leq \frac{C^2}{2} \sum_{i=1}^{N} \sum_{j=1}^{N} x_i x_j \tag{A17}$$

When $x_i$ and $x_j$ have the non-zero same dimension as $x_0$, we show that the masked inner product $x_i' x_j'$ is strictly less than $x_i x_j$. Suppose that $x_i^{(c)} = x_j^{(c)} = x_0^{(c)} = 1$, then $x_i x_j = \sum_{k \neq c} x_i^{(k)} x_j^{(k)} + 1 > \sum_{k \neq c} x_i^{(k)} x_j^{(k)} + 0 = x_i' x_j'$. In other cases, $x_i x_j = x_i' x_j'$. Thus, the mask technique reduces the upper bound determined by Equation (A17). □

Lemma 1.3 points out that the mask technique can lower the upper bound of regular terms of SVM, just like the CART models (even though the two are not for the same reason). With the same degree of regular term punishment, SVM with mask can get the same or better fitting results than SVM without mask. Therefore, we have the following corollary

**Corollary 1.2** *Compared with non-mask technique, for SVM with mask technique, the same or better results can be obtained.*

#### 1.4. Positional Encoding

Positional Encoding (PE) is usually added directly to input $x$ in order to add positional information.

**Definition 1.4** *The sin-cos PE used in Transformer of Transformer is* (Vaswani et al., 2017)

$$\begin{aligned} PE_{(pos,2i)} &= \sin\left(pos/10000^{2i/d_{\text{model}}}\right) \\ PE_{(pos,2i+1)} &= \cos\left(pos/10000^{2i/d_{\text{model}}}\right) \end{aligned} \tag{A18}$$

*where* $i$ *is the dimension of* $x$ $(i = 1, \cdots, d_{\text{model}})$ *and* $pos$ *is word's position* $(pos = 1, \cdots, n)$ *of sequence.*

#### 1.5. Ensemble Learning

Decision tree model uses ensemble learning, which can improve the generalization ability of the model and reduce the risk of overfitting. The theoretical basis is derived from Lemma 1.4 (Hoeffding, 1994)

**Lemma 1.4** (Hoeffding's Inequality) *Suppose* $X_i \in [a_i, b_i] (i = 1, \cdots, N)$ *is an independent random variable,* $\bar{X} = \frac{1}{N}\sum_{i=1}^{N} X_i$*, for* $\forall t > 0$*, we obtain*

$$\mathbb{P}(\left|\mathbb{E}[\bar{X}] - \bar{X}\right| \geq t) \leq 2\exp\left(-2N^2t^2 \Big/ \sum_{i=1}^{N}(b_i - a_i)^2\right) \tag{A19}$$

For convenience, we assume that the decision tree model consists of $T$ CARTs $f_k(k = 1, \cdots, T)$, each CART is generated by a subdataset of size $n$, Further assume that $f_k$ is independent of each other. The dataset is standardized, i.e., $\forall y_i \in (0, 1)$, by Lemma 1.4 we obtain

**Theorem 1.2** *For $T$-CARTs model and $\forall t > 0, \forall i > 0$*

$$\mathbb{P}(|y_i - \hat{y}_i| \geq t) = \mathbb{P}\left(\left|y_i - \frac{1}{T}\sum_{k=1}^{T} f_k(x_i)\right| \geq t\right) \leq 2\exp\left(-2T^2t^2\right) \tag{A20}$$

The upper bound of the error given in Theorem 1.2 decreases with the number of trees $T$, thus ensemble learning can effectively reduce the training error compared to a single CART.

#### 1.6. Learning Rate Adjustment Strategy

In our work, all optimizers use adaptive optimization algorithm *Adam* (Kingma and Ba, 2014). However, the adaptive optimization algorithm may converge to a poor local optimal solution (Keskar and Socher, 2017), so it is necessary to use a specific learning rate adjustment strategy for batter optimization. We use two different learning rate adjustment strategies, as shown in Figure A1.

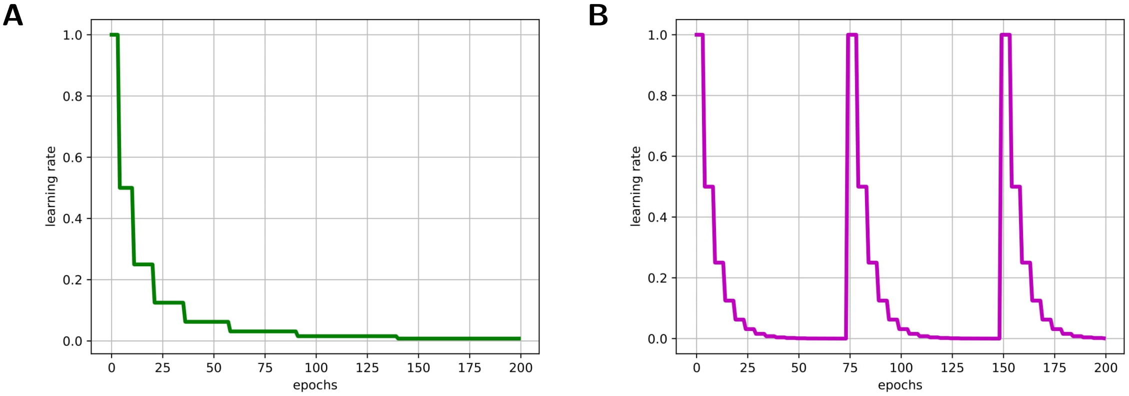


**Figure A1.** Learning rate adjustment strategies. (**A**) Step decay strategy. (**B**) Cyclic decay strategy.

**Definition 1.5** *Step decay strategy is a learning rate adjustment strategy in which the step size $s$ increases exponentially and the learning rate $l$ decreases exponentially. Suppose the decreasing rate of learning rate is $\alpha$ and the increasing rate of step size is $\beta$, the initial learning rate is $l_0$, initial step is $s_0$, then at step $T$, the learning rate is*

$$l = \alpha^n l_0 \quad \text{where } T \in \left[\frac{\beta^n - 1}{\beta - 1}s_0, \frac{\beta^{n+1} - 1}{\beta - 1}s_0\right) \tag{A21}$$

**Definition 1.6** *Cyclic decay strategy is a learning rate adjustment strategy in which the learning rate $l$ cycles by cycle size $c$ and decreases within cycle. Suppose that the decreasing step size within the cycle is $s$, the number of cyclic steps is $cs$, the decreasing rate of learning rate is $\alpha$, the initial learning rate is $l_0$, then the learning rate $l$ of the $T$ step is*

$$l = \alpha^{\lfloor t/s \rfloor} l_0 \quad \text{where } T \equiv t \mod cs \tag{A22}$$

In this work, LSTM uses step decay strategy, where $s_0 = 20$ and $\alpha = 0.10$. Transformer uses cyclic decay strategy, where $s = 10$, $c = 3$ and $\alpha = 0.10$.

#### 2. Details of Model Implementations

In the implementations of all models, the structure without EVMP is the same as that of BaseEncoder with EVMP. Therefore, this appendix only lists the details of EVMP implementations. For all models, the batch size is set to 16, the $L_2$ regularization coefficient of models using Adam is set to $10^{-5}$, both of which are not stated otherwise. In addition, for each input promoter shorter than 85 in length, we pad it with B (Blank, all-zero vector) at the end. In the following tables, we denote dropout ratio by *dp* and learning rate by *lr*.

##### 2.1. LSTM

The hyperparameters of EVMP-LSTM are shown in Table A1. Different from other models, the input of the VarEncoder of EVMP-LSTM is a *mutation number*. Each distinct mutant subsequence has a unique mutation number. Specifically, a mutation is generated at position $j$-th with respect to the $i$-th base promoter, while the $k$-mer mutant subsequence is one of the $m$-th of $2^k$ distinct mutant subsequences and can therefore be numbered by $(i, j, m)$. Each distinct number $(i, j, m)$ will be mapped to an embedding in $\mathbb{R}^{200}$. The detailed implementation of EVMP-LSTM is shown in Figure A2.

**Table A1.** Hyperparameters of EVMP-LSTM

| Model | input size | hidden units | seq len | dp | LSTM dp | LSTM layers | $k$ | lr |
|---|---|---|---|---|---|---|---|---|
| BaseEncoder | 5 | 1000 | 85 | 0.5 | 0.05 | 5 | — | $10^{-4}$ |
| VarEncoder | $1 \rightarrow 200$ | 200 | 10 | | | | 3 | |

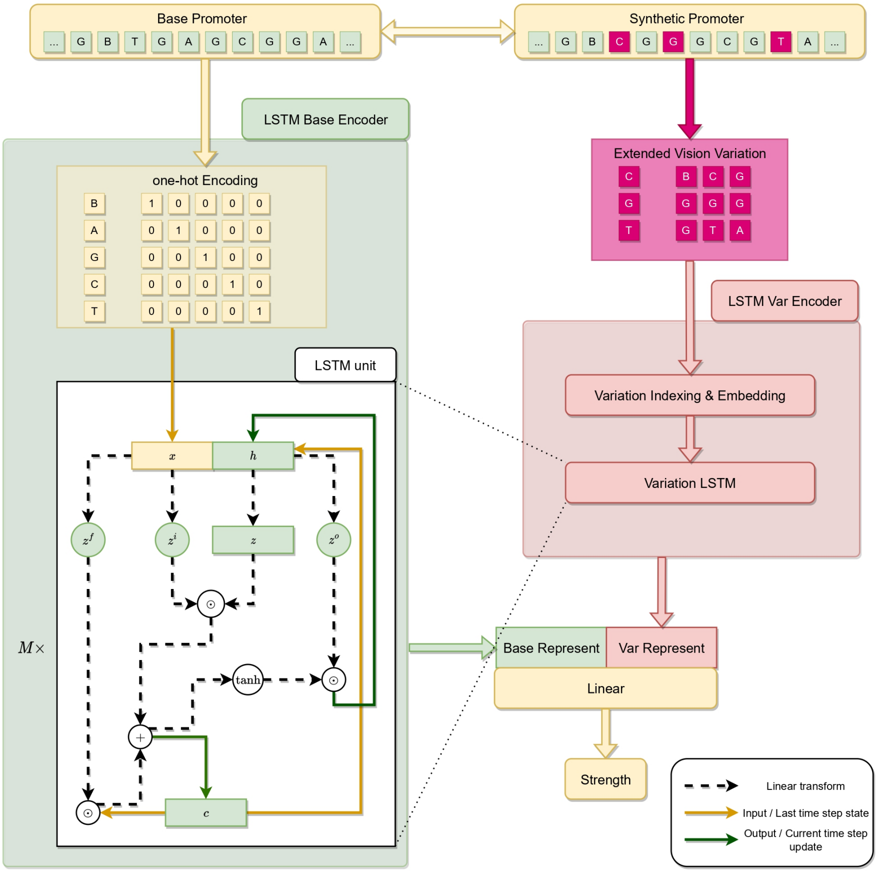


**Figure A2.** The implementation diagram of EVMP-LSTM.

#### 2.2. Transformer

The hyperparameters of EVMP-Transformer are shown in Table A2. The length of each base becomes 32 for BaseEncoder and 8 for VarEncoder, and the maximum number of mutations is set to 10. As mentioned in the main text, the BaseEncoder uses the Position Encoding of each base, while the VarEncoder uses the Position Encoding of each mutation position. The detailed implementation of EVMP-Transformer is shown in Figure A3.

**Table A2.** Hyperparameters of EVMP-Transformer

<table>
<tr><th>Model</th><th>word size</th><th>seq len</th><th>heads</th><th>FFN dim</th><th>$k$</th><th>PE</th><th>lr</th></tr>
<tr><td>BaseEncoder</td><td>5 → 32</td><td>85</td><td rowspan="2">8</td><td rowspan="2">2048</td><td>—</td><td>position based</td><td rowspan="2">0.0001</td></tr>
<tr><td>VarEncoder</td><td>5 → 8 × 10</td><td>10</td><td>5</td><td>mutation based</td></tr>
</table>

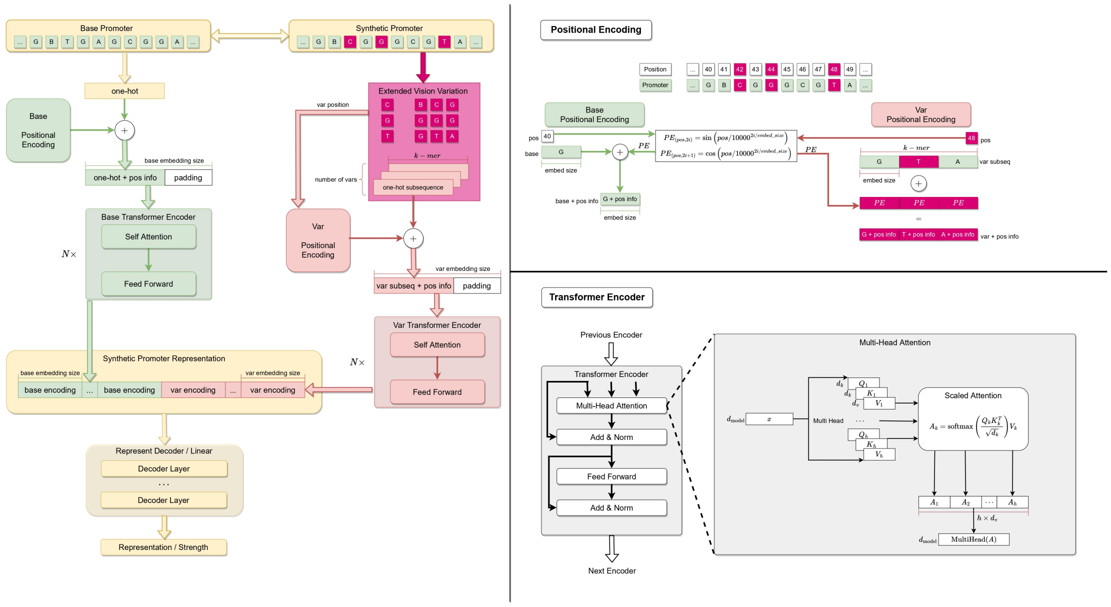


**Figure A3.** The implementation diagram of EVMP-Transformer.

#### 2.3. ML Models

The hyperparameters of ML models are shown in Table A3. The ML models with and without EVMP are exactly the same, except that the input of VarEncoder is masked synthetic promoter. In particular, XGBoost performs a grid search over the learning rate, the number of estimators and the maximum depth.

**Table A3.** Hyperparameters of ML models

<table>
<tr><th>Model</th><th>lr</th><th>regular $C$</th><th>estimators</th><th>max depth</th><th>folds</th><th>$k$</th></tr>
<tr><td>SVM</td><td></td><td>1</td><td></td><td></td><td></td><td rowspan="4">8</td></tr>
<tr><td>Random Forest</td><td></td><td></td><td>50</td><td></td><td></td></tr>
<tr><td>XGBoost</td><td>0.2, 0.1, 0.05</td><td></td><td>10, 50, 100, 300</td><td>3, 5, 7</td><td>5</td></tr>
<tr><td>GBDT</td><td>0.1</td><td></td><td>200</td><td>5</td><td></td></tr>
</table>

#### 2.4. Data Augmentation

Table A4 lists the first 10 base promoters used in the data augmentation experiments.

**Table A4.** Multiple base promoters used for data augmentation. The first 10 synthetic promoters were selected as the base promoters, and the mutation situation was basically consistent with the distribution of the dataset. $P_1$ is Trc promoter ($P_{\mathrm{trc}}$), $Ms$ is the number of mutations.

| ID | Base Promoter | Ms |
|---|---|---|
| $P_1$ | TTGACAATTAATCATCCGGCTCGTATAATGTBGTGGAATTGBTGAGCGGABTAACAATTTCACACAGGAAABCAGACC | 0 |
| $P_2$ | TTGACAATTAATCATCCGGCTCGTATAATGTBGCGGAATTGBTAAGCGGABTAACAATTTCACACAGGAAABCAGACC | 2 |
| $P_3$ | TTGACAATTAATCATCCGGCTCGTATAATGTBGTGGAATTGBTGGGCGGABTAACAATTTCACACAGGAAABCGGACC | 2 |
| $P_4$ | TTGACAATTAATCATCCGGCTCGTATAATGTBGTGGAATTGBCGGGCGTABTAACACTTTCACACAGGAAGBCAGACC | 5 |
| $P_5$ | TTGACAATTAATCACCCGGCTCGTATAATGTBGTGGAGTTGBTGAGCAGABTAACAATTTCACACAGGAAABCAGACC | 3 |
| $P_6$ | TTGACAATTAATCATCCGGCTCTTATAATGTBGTGGAATTGBTTATCTGCBTCACAATTCCACACAGGAAABCAGACC | 7 |
| $P_7$ | TTGACAATTAATCATCCGGCTCATATAGTGTBGTGGAATTGBTGGGCGGABTAACACTTTCACACAGGAAABCAGACC | 4 |
| $P_8$ | TTGACAATTAATCATCCGGCTCGTATAATGCBGTGGAATTGBTGGGCAGABTAGCACTTTCACACAGGAAABCAGACC | 5 |
| $P_9$ | CTGACAATTAATCATCCGGCTCGTATAATGTBGTGGAATTGBTGAGCAGABTAACAATTTCACACAGGAAABCAGACC | 2 |
| $P_{10}$ | TTGACAATTAATCATCCGGCTCTTATAATGTBGTGGAATTGBCGGGCGGABTAACAATTCCACACAGGAAABCAGACC | 4 |

L Breiman, JH Friedman, R Olshen, and CJ Stone. Classification and regression trees. 1984.

Tianqi Chen and Carlos Guestrin. Xgboost: A scalable tree boosting system. In *Proceedings of the 22nd acm sigkdd international conference on knowledge discovery and data mining*, pages 785–794, 2016.

Corinna Cortes and Vladimir Vapnik. Support-vector networks. *Machine learning*, 20(3):273–297, 1995.

Jerome H Friedman. Greedy function approximation: a gradient boosting machine. *Annals of statistics*, pages 1189–1232, 2001.

Wassily Hoeffding. Probability inequalities for sums of bounded random variables. In *The collected works of Wassily Hoeffding*, pages 409–426. Springer, 1994.

Nitish Shirish Keskar and Richard Socher. Improving generalization performance by switching from adam to sgd. *arXiv preprint arXiv:1712.07628*, 2017.

Diederik P Kingma and Jimmy Ba. Adam: A method for stochastic optimization. *arXiv preprint arXiv:1412.6980*, 2014.

Harold W Kuhn and Albert W Tucker. Nonlinear programming. In *Traces and emergence of nonlinear programming*, pages 247–258. Springer, 2014.

Sergios Theodoridis. *Machine learning: a Bayesian and optimization perspective*. Academic press, 2015.

Ashish Vaswani, Noam Shazeer, Niki Parmar, Jakob Uszkoreit, Llion Jones, Aidan N Gomez, Łukasz Kaiser, and Illia Polosukhin. Attention is all you need. *Advances in neural information processing systems*, 30, 2017.
